# Microfluidic Modulation of Microvasculature in Microdissected Tumors

**DOI:** 10.1101/2024.09.26.615278

**Authors:** Tran N. H. Nguyen, Lisa F. Horowitz, Brandon Nguyen, Ethan Lockhart, Songli Zhu, Taranjit S. Gujral, Albert Folch

## Abstract

The microvasculature within the tumor microenvironment (TME) plays an essential role in cancer signaling beyond nutrient delivery. However, it has been challenging to control the generation and/or maintenance of microvasculature in *ex vivo* systems, a critical step for establishing cancer models of high clinical biomimicry. There have been great successes in engineering tissues incorporating microvasculature *de novo* (*e.g.*, organoids and organs-on-chip), but these reconstituted tissues are formed with non-native cellular and molecular components that can skew certain outcomes such as drug efficacy. Microdissected tumors, on the other hand, show promise in preserving the TME, which is key for creating cancer models that can bridge the gap between bench and bedside. However, microdissected tumors are challenging to perfuse. Here, we developed a microfluidic platform that allows for perfusing the microvasculature of microdissected tumors. We demonstrate that, compared to diffusive transport, microfluidically perfused tissues feature larger and longer microvascular structures, with a better expression of CD31, a marker for endothelial cells, as analyzed by 3D imaging. This study also explores the effects of nitric oxide pathway-related drugs on endothelial cells, which are sensitive to shear stress and can activate endothelial nitric oxide synthase, producing nitric oxide. Our findings highlight the critical role of controlled perfusion and biochemical modulation in preserving tumor microvasculature, offering valuable insights for developing more effective cancer treatments.

## 1. INTRODUCTION

The fact that tumors are highly changing and rich ecosystems makes the engineering of *ex vivo* tumor models especially challenging. The tumor microenvironment (TME) is a very dynamic ensemble of proliferating tumor cells, migratory or rearranging stromal cells (such as immune and vascular cells as well as fibroblasts), extracellular matrix (ECM), and circulating fluids such as blood and lymph in their respective microvascular compartments or interstitial fluid.^1,2^ The abnormal structure^3–5^ and function^6,7^ of tumor vasculature hinder efficient nutrient and oxygen delivery while also causing an increase in interstitial fluid pressure (IFP), which drives abnormal interstitial flow within the TME.^8,9^ In normal tissues, fluid homeostasis is maintained through a balance of blood flow, lymphatic drainage, and interstitial flow, resulting in near-zero IFP.^10^ This balance is disrupted in tumors due to the disordered structure and impaired function of tumor vessels, including hyperpermeable blood vessels and solid stress-induced compression of both blood and lymphatic vessels.^11,12^ The excessive leakiness of tumor vessels, driven by defects in the endothelial lining and abnormal vessel walls, disrupts pressure gradients, causing IFP to rise to levels close to microvascular pressure.^13^ This elevated IFP, compounded by the lack of functional lymphatic vessels and the contractile activity of stromal elements, leads to abnormal interstitial flow, where fluid moves from high-pressure tumor regions into surrounding tissues.^5,14,15^ The steep IFP gradient at the tumor boundary generates an outward fluid flow that not only causes peritumor edema but also transports growth factors and cancer cells,^16^ promoting cancer cell migration and metastasis.^17^ The increased interstitial flow exerts mechanical stress on the ECM and stromal cells,^18^ facilitating matrix remodeling, fibroblast activation,^19^ and enhanced secretion of pro-tumorigenic factors such as TGF-β.^20^ Additionally, endothelial sprouting contributes to the disorganized and leaky vasculature,^21^ further exacerbating the alteration of normal fluid flow dynamics and altering angiogenic and lymphangiogenic processes.^9^ In sum, the interplay between interstitial flow and the tumor vasculature creates a complex landscape that is critical for understanding tumor progression and therapeutic outcomes. However, bioengineering both the microvasculature and interstitial flow conditions in *ex vivo* models has proven challenging, particularly in systems that maintain the native TME.

Microfluidic technology offers promising solutions to the critical need for accurate tumor models that seek to accurately replicate the TME with functional microvasculature. Bioengineered vascular structures enable precise control of vascular microenvironment cues, mimic interstitial flow, and allow for the perfusion of TME constructs *in vitro*.^22,23^ Functional assays on live intact tumor tissue, such as tumor slices^24–26^ or microdissected tumors (μDTs), also known as *ex vivo* tumor fragments,^27–32^ effectively preserve the TME and have significant potential for expanding drug testing but fail to retain the native vasculature. Recent studies have developed methods to support tumor explants with a vascularized bed, helping maintain tissue anatomy and viability by providing a dynamic interface with the surrounding environment. These support structures mimic some aspects of physiological conditions, such as nutrient and oxygen, but do not directly perfuse the internal vasculature of the tumor tissue.^33–35^ In contrast, perfusion systems developed for brain slices maintain interstitial flow, enhancing the viability and functional preservation of brain slices by providing a consistent supply of nutrients and oxygen and effectively removing waste.^36,37^ These non-tumor slice studies highlight the importance of perfusion in maintaining tissue functionality and offer insights into potential improvements for tumor explant models. Recently, we developed an *ex vivo* tumor fragment approach using regularly sized, cuboidal-shaped μDTs (“cuboids”) cut with a mechanical apparatus. This procedure can produce thousands of cuboids (∼400 μm^3^) from ∼1 cm³ of solid tumor in ∼30 min. These cuboids are never dissociated and retain much of the native TME, including immune cells and microvasculature, which enables the exploration of numerous drug treatments and therapies within TME-preserving tissue units.^38,39^ However, while μDTs retain significant portions of the TME, controlling the generation and/or maintenance of native vasculature in these tissues remains a challenge.^33^ Integrating dynamic flow systems, similar to those used in brain slice models, could help sustain vascular integrity and study interstitial flow effects in tumor tissues.

Several research groups have explored methods like orbital shaking,^40^ agitation,^41^ and perfusion bioreactors^42,43^ to improve μDT viability and integrity for extended periods across various cancer types. However, these studies have primarily focused on overall tissue health rather than specifically addressing the effects of fluid flow on microvasculature. The potential impact of such flow-based systems on the microvasculature within these 3D cultures remains underexplored, highlighting a critical gap in our understanding of how biophysical forces influence tumor vascularization *in vitro*. Given the central role of the tumor vasculature and the influence of interstitial flow on its formation and maintenance, a comprehensive understanding of these dynamics is crucial for developing effective therapeutic strategies. Here we found that, by establishing controlled, continuous fluid flow through mouse tumor cuboids with a microfluidic platform, we could support the structural integrity of 3D microvascular networks within the cuboids’ TME. Additionally, we investigated the effects of different flow magnitudes on microvascular integrity. We also examined the impact of vascular-disrupting agents (VDAs), such as Combretastatin A-4 (CA-4),^44^ and tested drugs affecting the nitric oxide (NO) pathway, including Nitroglycerin (NTG), a vasodilator,^45^ and nitric oxide synthase inhibitor, Nω-nitro-l-arginine (L-NNA).^46^ By analyzing how these factors impacted the microvascular structure and function of the vasculature within the TME, we aimed to identify potential therapeutic strategies that disrupt tumor blood supply and modulate the NO pathway, inhibiting tumor growth and enhancing therapeutic outcomes.

## 2. RESULTS

### 2.1. Experimental Process Overview

The experimental workflow, schematized in **Fig. 1A**, encompassed from tumor generation and microdissection to cuboid perfusion and data analysis. First, we generated xenograft tumors through injections in mice. We then processed the tumors into uniform cuboids, ensuring consistency in size and shape for subsequent experiments according to a previously described protocol.^32,39^ Next, we loaded the cuboids into the microfluidic device and perfuse them with culture medium, comparing these to no-flow culture (*i.e.*, diffusive transport) conditions. In some experiments, we exposed the cuboids to perfusion and no-flow conditions with 70 kDa dextran Texas Red. At the end of the perfusion, to visualize the microvasculature we immunostained the cuboids with an antibody to CD31, a marker for endothelial cells, and employed tissue-clearing techniques to enhance tissue transparency for confocal imaging. Using high-resolution 3D confocal microscopy, we imaged the cuboids directly within the microfluidic device, allowing for detailed visualization of the microvasculature architecture. We captured z-stack images during the confocal microscopy session, enabling 3D reconstruction and comprehensive analysis with Imaris software. This analysis provided quantitative data on various parameters, including microvasculature volume, branching, and overall structural integrity.

**Figure 1.**
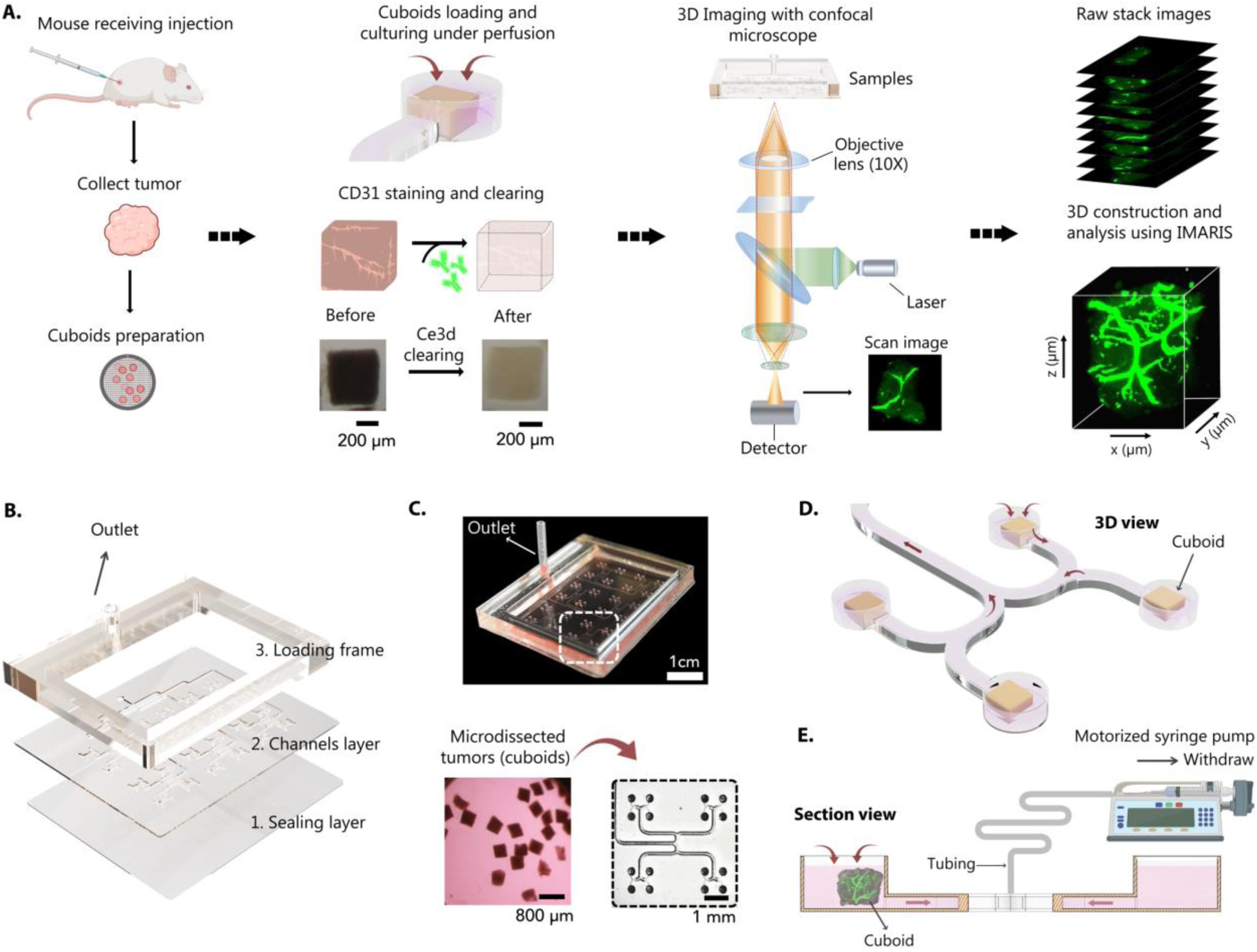
Overview of microfluidic device development and experimental workflow: **A.** Diagram of the Experimental Workflow: generation of xenograft tumors via injection in a mouse and subsequent processing of the tumor into cuboids; Loading of cuboids into the device for perfusion, followed by CD31 immunostaining and tissue clearing; 3D confocal microscopy imaging performed directly on the cuboids within the device; Capture of z-stack images for 3D reconstruction and analysis with Imaris. **B.** Schematic representation detailing the microfluidic platform’s layered structure, comprising the sealing layer, channels layer, and loading frame, arranged from bottom to top. **C.** Photograph of the microfluidic device: The top panel shows the fully assembled device ready for cuboid culture in the medium; the bottom left depicts microdissected tumor tissues (cuboids); the bottom right focuses on a close-up section of the device displaying cuboids situated in four individual traps. **D.** A 3D illustration of four cuboids in their traps and the direction of flow inside the device. **E.** Cross-sectional schematic of the device connected to a motorized syringe pump.

### 2.2. Design and fabrication of the microfluidic device and fluidic control

The microfluidic device, designed to optimize tissue culture and imaging, features a layered assembly that facilitates fluid control. **Fig. 1B** illustrates the device’s layered structure, which includes a sealing layer, channel layer, and loading frame with an outlet arranged from bottom to top. This design enables efficient trapping of cuboids into addressable traps for culture. We designed the device with 96 hydrodynamic traps clustered in groups of four (**Fig. 1C**). This multi-stage design includes a loading and culture stage. The topmost layer, the loading frame, creates a fluid reservoir to hold the cuboid suspension. The microchannel layer below drains the standing solution into the traps, with 100 µm openings to prevent the ∼400 µm cuboids from entering the fluidic network. The microchannel layer features a network of bifurcating channels (∼400 µm width and ∼350 µm in depth), evenly propagating suction from an external pressure source to each trap, enhancing tissue capture efficiency. Larger connecting channels reduce fluidic resistance, maximizing the suction pull. **Fig. 1D** illustrates how the flow is used and controlled by a motorized syringe pump to trap the cuboids.

We fabricated the device using PMMA because it is inexpensive, optically clear, and biocompatible. Unlike polydimethylsiloxane (PDMS), PMMA does not absorb drugs easily, making it suitable for drug testing applications.^47,48^ The PMMA device is also amenable to rapid prototyping via CO_2_ laser machining, enhancing cost-effectiveness, precision, and throughput. Moreover, the transparency of PMMA facilitates an on-chip tissue clearing protocol, enabling accurate 3D confocal imaging directly on the device (containing up to 96 isolated cuboids) and quantification of the microvasculature.^48^ To investigate flow mechanisms in maintaining 3D network formation, we used the device to control flow magnitude precisely with motorized syringe pumps (**Fig. 1E**). This setup enabled us to establish media perfusion through the entire network, confining tissues to restricted regions and regulating microenvironmental conditions.

### 2.3. Dextran perfusion analysis

To rule out diffusive transport, we measured the penetration of a large MW dye within the cuboids by perfusing the microfluidic devices with cultured medium containing dextran Texas Red 70 kDa under different flow conditions. Since testing all flow conditions at once wasn’t feasible, we divided the experiments across multiple tumors to assess the effects of various flow conditions over time. **Fig. 2A** presents the experimental design and timelines for different flow conditions at 2 hrs and 24 hrs. After perfusion, we cleared the cuboids and performed confocal imaging to obtain the red fluorescence intensity in all the interior points of the cuboids. **Fig. 2B** first illustrates the confocal imaging presentation, showing the top panel with orthogonal slices at a 100 µm depth and the bottom panel with z-axis maximum intensity projections. **Fig. 2C, E** present the confocal imaging results for 2 hrs and 24 hrs under different flow conditions (0.05 mL/hr, 0.1 mL/hr, 0.15 mL/hr), a control condition with no flow, and the orbital shaking condition. The confocal images revealed that cuboids subjected to flow exhibited higher dextran intensity compared to the no-flow control, and dextran intensity was higher in cuboids perfused for 24 hrs than only 2 hrs, indicating enhanced penetration under flow conditions. The dextran intensity average, calculated by averaging the red-fluorescence intensity over the entire 3D volume of the cuboids (**Fig. 2D, F**), confirmed that dextran intensity under all flow conditions was significantly higher than that of the no-flow (diffusion) control. Statistical analyses showed significant differences particularly between 0.1 mL/hr and 0.15 mL/hr at both 2 hrs and 24 hrs. The results further emphasize that the flow rate plays a critical role in enhancing dextran penetration within the cuboids, particularly at higher flow rates, suggesting that optimizing flow conditions could significantly improve delivery efficiency in microdissected tumor cuboid models.

**Figure 2.**
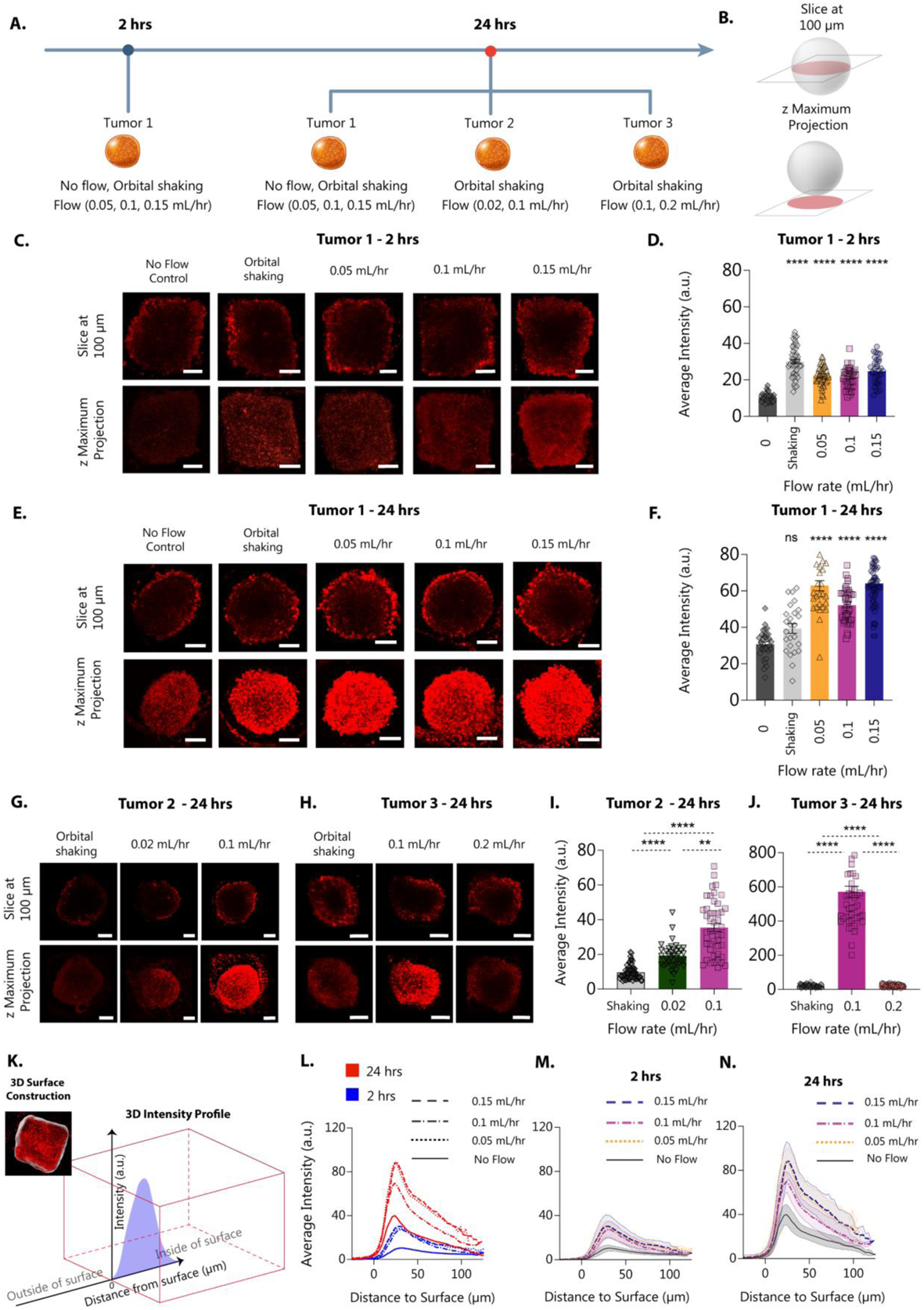
Dextran perfusion and quantitative analysis of cuboids in the microfluidic device. **A.** Flow chart illustrating the experimental design and timelines for the 2 hrs and 24 hrs conditions. **B.** Conceptual drawing showing the confocal imaging presentation. The top panel depicts orthogonal slices at a 100 µm depth of the cuboid for each condition, providing a cross-sectional view. The bottom panel displays z-axis maximum intensity projections for all conditions. **C.** Confocal visualization of dextran perfusion for cuboids from tumor 1: sequential confocal images of dextran Texas Red 70 kDa perfusion under various conditions at 2 hrs. Scale bar, 100 µm. **D.** Quantification of dextran intensity for cuboids from tumor 1: plots showing the average fluorescence intensity of dextran Texas Red at 2 hrs under different conditions. **E.** Confocal visualization of dextran perfusion for cuboids from tumor 1 at 24 hrs under all conditions. **F.** Quantification of dextran intensity at 24 hrs for cuboids from tumor 1 under different flow and control conditions. **G-H.** Confocal visualization of dextran perfusion for additional flow rates in cuboids from tumor 2 and tumor 3 at 24 hrs. **I-J.** Quantification of dextran intensity at 24 hrs for the additional flow conditions for cuboids from tumors 2 and 3. Data represent mean ± S.E.M. for approximately 30-50 cuboids per condition. Statistical significance was determined using ordinary one-way ANOVA with Tukey’s post-test for 2 hrs, and nonparametric Kruskal-Wallis one-way ANOVA with Dunn’s post-test for 24 hrs, with significance denoted as *p<0.05, ***p<0.001, and ****p<0.0001; non-significant findings are indicated by ‘ns’ for p>0.05. **K.** 3D surface reconstruction and intensity profiling: 3D reconstruction of the cuboid surfaces generated in Imaris, with a schematic detailing how dextran intensity profiles were calculated based on the fluorescence signal variation relative to the distance from the surface. **L.** Comparative intensity profiles: an overview of dextran intensity profiles across all conditions and time points for cuboids from tumor 1. **M.** Detailed intensity profile for the 2 hrs condition: expanded graph showing the dextran intensity distribution at 2 hrs, with shading representing standard deviation. **N.** Detailed intensity profile for the 24 hrs condition: expanded graph showing the dextran intensity distribution at 24 hrs, with shading representing standard deviation. Data represent approximately 30-50 cuboids per condition, with some variation in group sizes.

We also investigated whether orbital shaking could enhance dextran penetration. At 2 hours, the average intensity under orbital shaking conditions was comparable to that of flow (**Fig. 2C, D**). However, the intensity dropped significantly compared to all flow conditions at 24 hrs (**Fig. 2E, F**). While cuboids subjected to orbital shaking exhibited slightly higher dextran intensity than the no-flow control, the increase was not statistically significant at 24 hrs. In contrast, cuboids subjected to flow conditions demonstrated significantly higher dextran intensity than both no-flow and orbital shaking conditions, indicating that while orbital shaking may offer some improvement, fluid flow remains the more effective method for enhancing dextran penetration within the cuboids. These findings emphasize the advantage of controlled fluid flow over orbital shaking in promoting effective molecular delivery in microdissected tissue models.

We also sought to examine whether cuboid volume, alongside flow conditions, could affect dextran penetration. To assess the impact of cuboid volume on dextran penetration, we analyzed correlations between cuboid volume and normalized dextran intensity under different flow rates and a no-flow control at 2 hrs and 24 hrs (**Fig. S**1). The only strong correlations at both 2 hrs and 24 hrs are observed for cuboids perfused at the lowest flow rate (0.05 mL/hr), whereas cuboids with twice as much volume had ∼33% and ∼31% less dextran intensity, respectively. At higher flow rates and under no-flow control, no significant correlations were observed, indicating that the effect of cuboid volume was negligible at these conditions. At 24 hrs, weaker correlations were observed for the no-flow condition and the 0.15 mL/h flow rate, where cuboids with twice the volume showed ∼20% and ∼25% less dextran intensity, respectively (**Fig. S1**B). The slopes for the 0.05 mL/h flow condition at both time points were statistically significant, suggesting that flow rate plays a critical role in the relationship between cuboid volume and dextran penetration, with the effect being more pronounced at lower flow rates. These findings suggest potential diffusion limitations or volumetric dilution effects, where larger cuboids may retain less dextran due to distribution within a larger space, especially at lower flow rates where diffusion dominates.

Considering that there is a variability in cuboid sizes due to the cutting process and that the volume of cuboids can influence the intensity of dextran, we conducted additional analyses to address this variability. **Fig. S2** examines whether the results presented in **Fig. 2D, F** still hold when we select a range of cuboid volumes where no correlation between volume and dextran intensity is observed. **Fig S2** shows results from 2 hrs and 24 hrs conditions. **Fig. S2**A, D illustrates the relationship between cuboid volume and average dextran intensity for all cuboids across various flow conditions. A dashed box highlights the cut-off volume range where no significant correlation was found between cuboid volume and dextran intensity. This range was selected to control for variability in cuboid size, ensuring that the influence of volume on dextran intensity was minimal. **Fig. S2**B, E zoom in on cuboids within this cut-off volume range. **Fig. S2**C, F show the average dextran intensity for flow rates of 0.05, 0.1, and 0.15 mL/hr compared to the no-flow condition and orbital shaking, specifically for cuboids within the cut-off volume range. The data indicate that even when controlling for cuboid volume variability, significant differences in dextran intensity remain under different flow rates. This analysis supports the conclusion that the observed differences in dextran intensity are primarily due to the applied flow rates rather than variations in cuboid volume. Therefore, the results of **Fig. 2D, F** are robust, even after accounting for size-related variability in cuboid volume.

To further explore the limits of dextran perfusion, we tested additional flow rates and compared them to orbital shaking, which previously showed slightly higher dextran penetration than no-flow conditions. This analysis aimed to identify the flow rate range that maximizes dextran penetration while avoiding the inefficiencies observed at extreme flow rates. As shown in **Fig. 2G**, confocal visualization at 24 hrs under flow rates of 0.02 mL/hr and 0.1 mL/hr, alongside orbital shaking, reveals that 0.1 mL/hr resulted in greater dextran penetration. The graph of dextran intensity (**Fig. 2I**) further confirms that the average dextran intensity is significantly lower at 0.02 mL/hr than at 0.1 mL/hr, though still higher than under orbital shaking. Similarly, the confocal images of dextran perfusion at 0.1 mL/hr, 0.2 mL/hr, and under orbital shaking shown in **Fig. 2H** demonstrate that 0.1 mL/hr again results in the highest dextran intensity (see graph in **Fig. 2J**). Notably, the average dextran intensity at 0.1 mL/hr is significantly higher than at 0.2 mL/hr and orbital shaking, with no significant difference observed between 0.2 mL/hr and orbital shaking. These results indicate that flow rates ranging from 0.05 mL/hr to 0.15 mL/hr are optimal for dextran perfusion, with no significant differences observed among them. In contrast, flow rates lower than 0.05 mL/hr (such as 0.02 mL/hr) and higher than 0.15 mL/hr (such as 0.2 mL/hr) are less effective. The result highlights the importance of maintaining flow rates within the 0.05 mL/hr to 0.15 mL/hr range to maximize dextran penetration compared to no-flow and orbital shaking conditions.

Measurements of dextran distribution within the cuboids provided further evidence of the microfluidic device’s efficacy in enhancing molecular transport. 3D intensity profiles of dextran within the cuboids (**Fig. 2K**) demonstrated how fluorescence intensity decreases as the distance from the surface increases, indicating the extent of dextran penetration into the cuboids. A comparison of dextran intensity within 50 µm of the surface at different time points and flow conditions revealed that dextran penetration is significantly enhanced over time and with increased fluid flow (**Fig. 2L-N**). These results confirm that the microfluidic device facilitates effective perfusion and notably improves the delivery and distribution of molecules within 3D tissue models, particularly over extended perfusion periods.

### 2.4. Vasculature analysis within cuboids via CD31 confocal imaging

We hypothesized that externally applied flow to our cuboids would influence the growth or preservation of microvasculature in the TME of the cuboids, which would then be visible with a vascular marker (*i.e.*, CD31) and could be analyzed with various morphological metrics. To evaluate the effects of different flow rates on microvascular structures, we conducted a detailed vasculature analysis within cuboids using CD31 confocal imaging. The vascular analysis workflow in Imaris (**Fig. 3A**) illustrates the step-by-step process from a 3D confocal microscope image of a cuboid to generating CD31 immunostained surfaces and detailed “filament tracing” using the generated surfaces as masks for precise mapping. We used the “filament tracing” function in Imaris to collect detailed data on the microvascular architecture, including parameters such as branch length, branch average diameter, and overall network connectivity. Given that testing all conditions simultaneously wasn’t feasible, we broke up the experiment and used different tumors across multiple experimental setups (**Fig. 2A**). As before, we combined flow conditions (0.05, 0.1, and 0.15 mL/hr) with no-flow controls and orbital shaking to investigate their effects on vasculature. The analysis of the confocal images of CD31-stained cuboids for various flow rates and at different time points revealed notable trends in microvascular morphology. CD31 staining was consistently better maintained under flow conditions (0.05, 0.1, and 0.15 mL/hr) compared to no-flow controls at various time points (2 hrs, 24 hrs, and 72 hrs) (**Fig. 3B**). The presented confocal images were selected to represent an average cuboid for each condition. In general, all flow conditions showed better support for the vasculature compared to no-flow control. However, the highest (0.15 mL/hr) flow rate showed the most disconnections in the vasculature. These results suggest that, while flow is generally beneficial for either generating new vessels or maintaining vascular integrity (our terminal CD31 staining cannot distinguish between these two scenarios), excessively high flow rates might have detrimental effects. Additionally, we included orbital shaking conditions to validate the impact of different flow rates on microvascular structures. As shown in **Fig. 3B**, confocal images of cuboids subjected to orbital shaking at 2 hrs show better CD31 staining compared to the no-flow control, with a similar effect to that observed under the 0.15 mL/hr flow condition. However, at 24 hrs and 72 hrs, CD31 staining was better maintained under flow conditions compared to both no-flow and orbital shaking. Notably, the CD31 staining under orbital shaking was similar to the no-flow control, with both showing diminished staining relative to the flow conditions. This result highlights the importance of controlled perfusion over orbital shaking in maintaining vascular structures within the cuboids.

**Figure 3.**
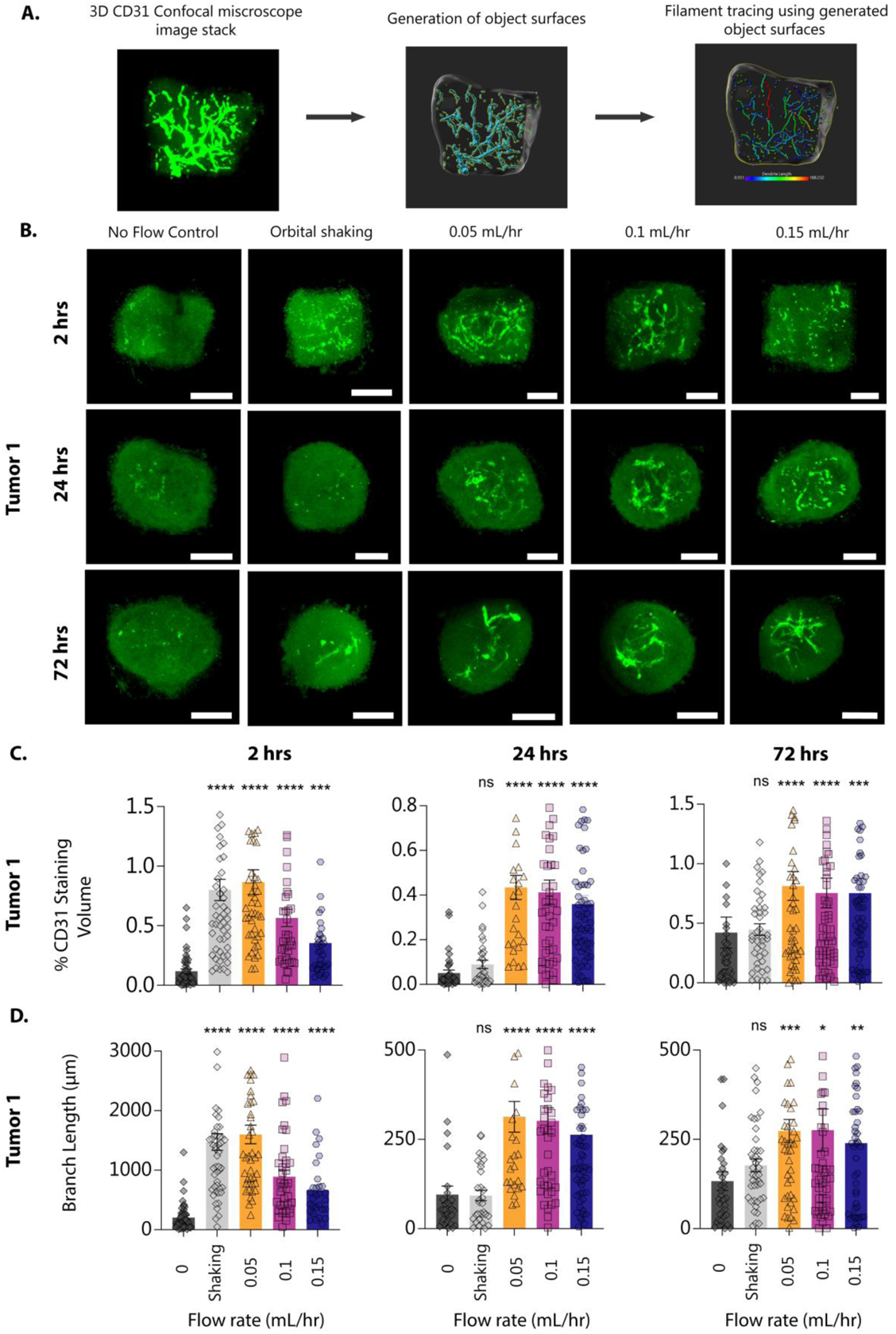
Vasculature analysis within cuboids via CD31 confocal imaging. **A.** Vascular analysis workflow in Imaris: The schematic illustrates the step-by-step process within Imaris, starting from a 3D confocal microscope image of a cuboid and progressing through the stages of generating CD31 immunostained surfaces to detailed filament tracing using the generated surfaces as masks for precise mapping. **B.** Confocal images of cuboids stained with CD31 being subjected to different flow rates (0.05, 0.1, and 0.15 mL/hr), no-flow control, and orbital shaking for different time points at 2 hrs, 24 hrs, and 72 hrs. Scale bar, 100 µm **C.** Percentage volume of CD31 staining. **D.** Branch length of all conditions over 2 hrs, 24 hrs, and 72 hrs. Data are represented as mean ± S.E.M. for 30-50 cuboids per condition. Statistical significance was determined using nonparametric Kruskal-Wallis one-way ANOVA with Dunn’s post-test, with significance denoted as *p<0.05, ***p<0.001, and ***p<0.0001; non-significant findings are indicated by ‘ns’ for p>0.05.

Quantitative analysis of CD31 staining volume (**Fig. 3C**) further supports these observations, with cuboids under flow conditions exhibiting significantly higher CD31 staining volume than those under no-flow control conditions. Since cuboid size changes over the culture period, we normalized the CD31 staining volume data to each cuboid’s outer volume. The percentage of CD31 staining volume normalized to the outer surface volume of the cuboids was highest under flow conditions (**Fig. 3C**). The 0.05 mL/hr flow rate resulted in the highest percentage of staining volume, followed by 0.1 mL/hr and 0.15 mL/hr, across 2 hrs, 24 hrs, and 72 hrs. While all flow rates significantly improved vascular maintenance compared to the no-flow control (as shown in the figure), the statistical tests between different flow conditions were not significant, indicating that all three flow rates are effective for maintaining vascular surfaces. Similarly, for orbital shaking, the percentage of CD31 staining volume at 2 hrs was comparable to the flow conditions (**Fig. 3C**). However, by 24 hrs and 72 hrs, the percentage of CD31 staining volume dropped significantly, reaching levels similar to the no-flow control. While orbital shaking may initially support vascular integrity, it fails to maintain it over extended periods compared to controlled perfusion. These findings align with our earlier observations on dextran penetration, where higher perfusion rates (0.05, 0.1, and 0.15 mL/hr) more effectively sustained vascular structures, whereas orbital shaking did not provide the same benefit.

Additionally, to assess the impact of cuboid volume on the percentage of CD31 staining volume, we analyzed the correlation between cuboid outer volume and the CD31 staining volume under various flow conditions and time points (2 hrs, 24 hrs, and 72 hrs). At 2 hrs (**Fig. S3**A) and 24 hrs (**Fig. S3**B), no significant correlations were observed between cuboid volume and CD31 staining volume across all conditions, indicating that the impact of cuboid volume on vascularization is minimal during the early stages of culture. By 72 hrs (**Fig. S3**C), however, the correlation between cuboid volume and the percentage of CD31 staining volume became more apparent, particularly under higher flow rates. At 72 hrs, cuboids with twice the volume showed approximately ∼28.55% less CD31 staining volume at 0.05 mL/h, ∼28% less at 0.1 mL/h, and ∼57% less at 0.15 mL/h, indicating a significant relationship between volume and vascularization, with the slopes for these conditions being statistically significant and highlighting the pronounced effect of flow rate in influencing the relationship between cuboid size and vascularization over time. The results highlight the importance of considering cuboid outer volume in vascularization studies, particularly when higher perfusion rates are applied over extended culture periods. Normalizing the CD31 staining volume to the outer surface volume of the cuboids is crucial for accurately assessing these effects, as it accounts for variations in cuboid size that could otherwise confound the results. The smaller cuboids consistently exhibited a higher percentage of CD31 staining volume, indicating more efficient vascularization at these flow rates.

We also explored many other metrics (available in Imaris) to evaluate morphological changes in the vasculature over time for the various flow rates. Since the vasculature appears to “disintegrate”, we evaluated measures of connectivity such as branching. The analysis of branch length (**Fig. 3D**) revealed that cuboids under flow conditions had significantly longer branch lengths compared to no-flow controls. The results show that the effect was consistent across all time points, with the 0.05 mL/hr flow rate showing the longest branch lengths at 2 hrs, though by 24 and 72 hrs, the 0.05 mL/hr and 0.1 mL/hr flow rates showed similar branch lengths, with 0.15 mL/hr resulting in shorter branches. Since some of these differences were not statistically significant, we conclude that some metrics might require more experimental confirmation.

The vasculature analysis within cuboids using CD31 confocal imaging revealed that flow conditions significantly improved the maintenance of microvascular structures compared to no-flow conditions. Flow rates of 0.05, 0.1, and 0.15 mL/hr all resulted in better preservation of CD31 staining, with higher average intensity and a greater percentage of CD31 staining volume. While the 0.05 mL/hr flow rate generally showed the highest performance across various metrics, including the percentage of staining volume and branch length, the differences between the flow rates were not statistically significant. Additionally, flow conditions promoted longer branch lengths within the vasculature, indicating better network maintenance. These findings suggest that implementing flow rates between 0.05 and 0.15 mL/hr can effectively sustain the microvascular architecture within cuboids, providing a robust environment for further biological studies.

Furthermore, we extended our CD31 staining analysis to include additional flow rates and orbital shaking to validate our previous findings. As shown in **Fig. S4**A, confocal images of cuboids subjected to flow rates of 0.02 and 0.1 mL/hr, alongside orbital shaking, again demonstrated better maintenance of CD31 staining under flow conditions, with orbital shaking leading to a marked reduction in staining at 24 hrs. The percentage of CD31 staining volume (**Fig. 4B**) were significantly higher under 0.1 mL/hr flow compared to 0.02 mL/hr and orbital shaking, further validating the critical role of flow in maintaining vascular integrity. Lastly, as depicted in **Fig. 4C**, cuboids exposed to flow rates of 0.1 and 0.2 mL/hr compared to orbital shaking showed that higher flow rates at 0.2 mL/h could lead to more disconnected vasculature than 0.1 mL/h. The analysis of percentage of CD31 staining volume (**Fig. 4D**) confirmed that 0.1 mL/hr provided superior vascular preservation compared to 0.2 mL/hr and orbital shaking, with statistically significant differences. By combining the findings from **Fig.4**, we confirm that maintaining a flow rate within the 0.05 mL/hr to 0.15 mL/hr range is optimal for preserving vascular structures within cuboids, as previously established in **Fig. 3**. Flow rates outside this range, as well as orbital shaking, result in reduced CD31 staining volume, underscoring the importance of controlled perfusion for maintaining microvascular integrity in cultured tissues.

**Figure 4.**
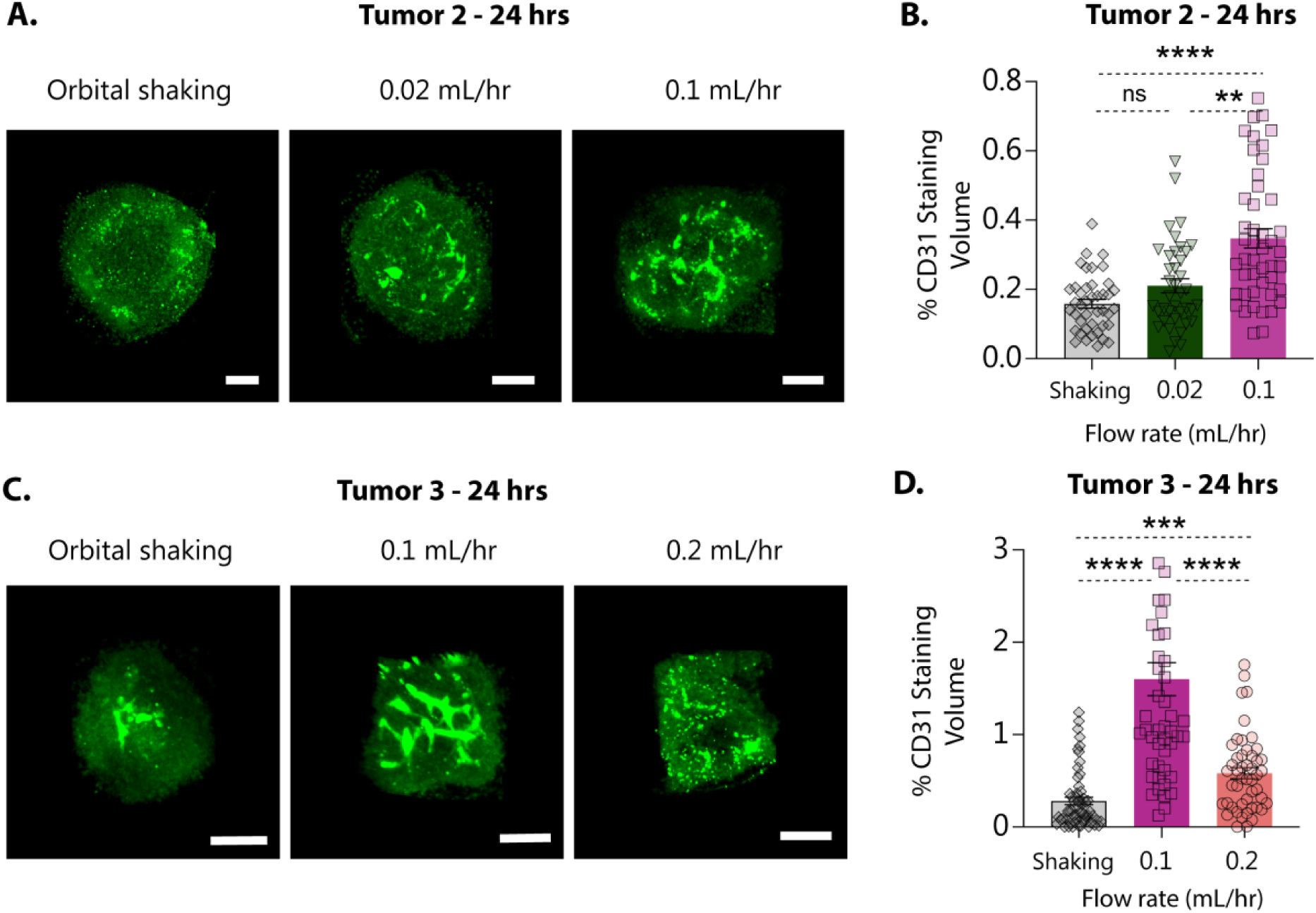
Vasculature analysis within cuboids via CD31 confocal imaging with different flow rates. **A.** Confocal images of cuboids from tumor 2 stained with CD31 subjected to different flow rates (0.02, 0.1 mL/hr) and orbital shaking for 24 hrs. Scale bar, 100 µm. **B.** Percentage volume of CD31 staining at 24 hrs. **C.** Confocal images of cuboids from tumor 3 stained with CD31 subjected to different flow rates (0.1, 0.2 mL/hr) and orbital shaking for 24 hrs. Scale bar, 100 µm. **D.** Percentage volume of CD31 staining at 24 hrs. Data are represented as mean ± S.E.M. for 30-50 cuboids per condition. Statistical significance was determined using nonparametric Kruskal-Wallis one-way ANOVA with Dunn’s post-test, with significance denoted as *p<0.05, ***p<0.001, and ***p<0.0001; non-significant findings are indicated by ‘ns’ for p>0.05.

### 2.5. Drug treatments modulate vascular characteristics in microfluidic cultures of cuboids

One of the central mechanisms influenced by controlled flow is the activation of endothelial nitric oxide synthase (eNOS), which leads to NO production, a critical mediator of vascular health.^49–51^ NO promotes vasodilation, enhancing blood flow, nutrient delivery, and waste removal, all of which are essential for supporting a healthy microvascular environment.^58,72^ Due to the disorganized structure, disrupted endothelial layers, and abnormal basement membrane support typically observed in tumor vasculature, flow has a vital influence in tumor physiology, including a key role in eNOS and NO release under shear stress (**Fig. 5A**).^52^ Different NO levels can either promote normal physiological processes or enhance tumorigenic and antitumor effects (**Fig. 5B**).^53,54^ Thus, we investigated the effects of various vascular-targeting drugs, such as L-NNA and CA-4, and tested drugs affecting the nitric oxide (NO) pathway, including Nitroglycerin (NTG), a vasodilator on the vascular characteristics of cuboids cultured in our microfluidic platform. L-NNA, an eNOS inhibitor, blocks NO production, leading to complex effects on vascularization. Inhibiting NO production causes a reduction in vasodilation, increasing vascular tone and resistance. This effect can increase vascular volume due to the reduced ability of blood vessels to dilate, contributing to higher vascular resistance and potential issues with blood flow and tissue perfusion, but decreases overall vascularization by inhibiting NO’s angiogenic effects.^51,55,56^ In contrast, CA-4, a tubulin-binding agent, targets and destabilizes microtubules in endothelial cells.

**Figure 5.**
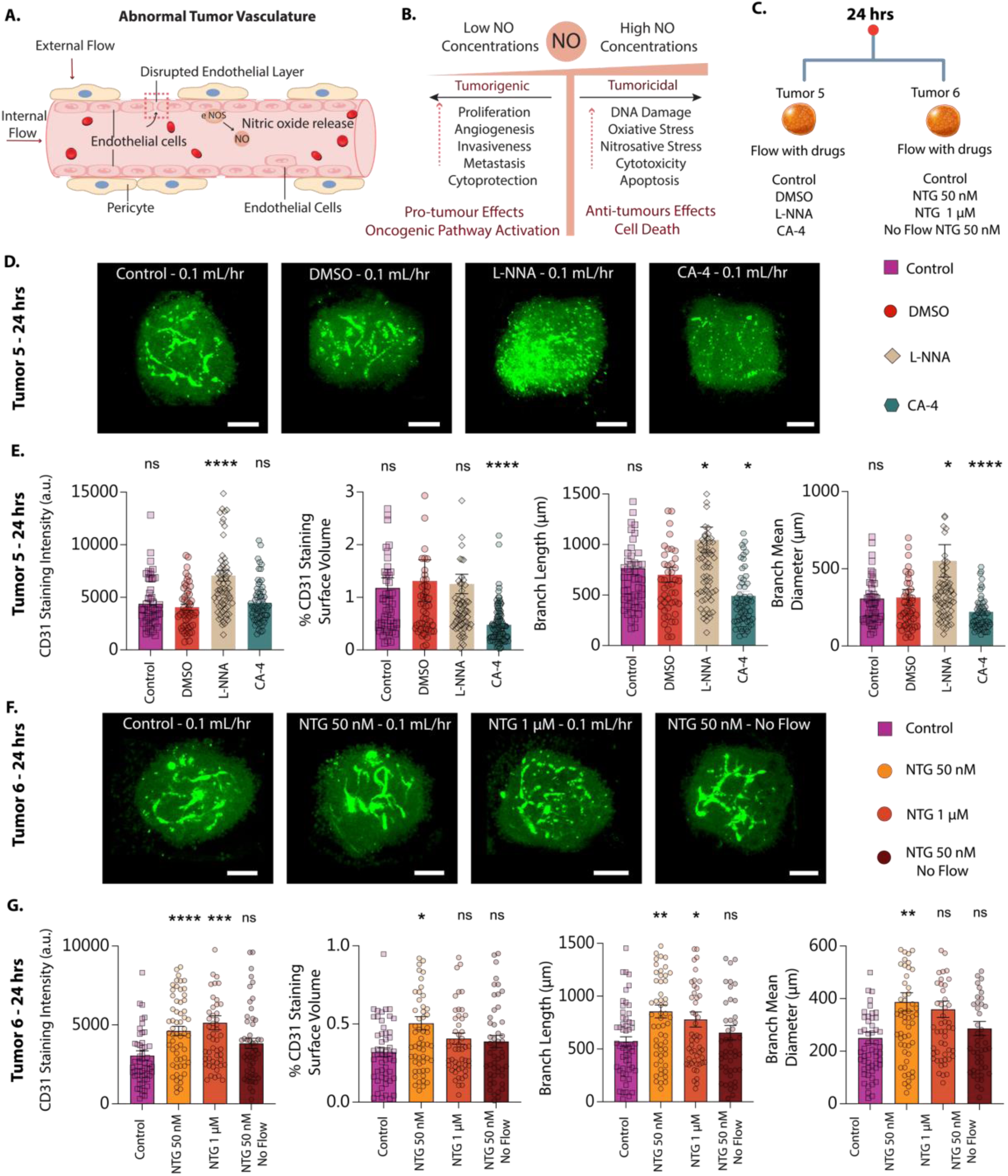
Drug treatment effects on microfluidic-cultured cuboids. A. Schematic representation of the effect of flow on tumor vasculature. Tumor vasculature is frequently structurally disorganized, with disrupted endothelial layers, poor pericyte coverage, and abnormal basement membrane support. The schematic also indicates the presence of eNOS on the endothelial cells and the release of NO, typically due to shear stress. **B.** Concentration-dependent effects of NO in cancer. The panel illustrates how different NO concentrations can have varying effects on cancer. Low NO levels can improve molecular processes that maintain normal physiology but may also promote tumorigenic, pro-tumor effects such as oncogenic pathway activation, increased proliferation, and angiogenesis. High NO levels, however, promote tumoricidal and antitumor effects, including DNA damage, oxidative/nitrosative stress, cytotoxicity, and apoptosis. **C.** Flow chart illustrating the experimental design of cuboids from tumors 5 and 6 under different drug treatments. **D.** Confocal imaging post-treatment presents CD31 immunostained cuboids after exposure to L-NNA, CA-4, or control conditions, including DMSO vehicle and normal flow without drug treatment. The images captured these conditions after 24 hrs of continuous flow at 0.1 mL/hr. Scale bar, 100 µm. **E.** Quantitative vascular analysis provides data from CD31 immunostained surfaces. This data illustrates the average intensity of CD31 staining, the percentage volume of CD31 staining, branch length, and branch average diameter categorized by treatment groups of all conditions over 24 hrs. **F.** Visualization of NTG treatment displays confocal microscopy images of CD31 immunostained cuboids. These cuboids were treated with NTG at 50 nM and 1 µM concentrations under flow conditions set to 0.1 mL/hr alongside a control group under the same flow without the drug and a static group treated with 50 nM NTG. Scale bar, 100 µm. **G.** Post-NTG quantitative vascular analysis displays analytical data on the average intensity of CD31 staining, the percentage volume of CD31 staining, branch length, and branch average diameter across all NTG treatment groups compared to controls. Statistical significance was determined using one-way ANOVA with the Kruskal-Wallis and Dunn’s post-test. Significance is indicated by *p<0.05, ***p<0.001, and ****p<0.0001, while ‘ns’ denotes non-significant differences for p>0.05.

Due to the complexity of testing all drug treatments in one experiment, we divided the study into multiple experiments across different tumors. **Fig. 5C** presents the experimental design for cuboids from tumors 5 and 6 under different drug treatments, with controlled flow rates and conditions across each experiment to analyze the vascular response.

We used confocal imaging to assess the impact of drug treatments on vascular structures within the cuboids after 24 hrs of continuous flow at 0.1 mL/hr (**Fig. 5D**). The images clearly show that L-NNA treatment enhances CD31 staining intensity and clusters vasculatures, while CA-4 treatment significantly reduces CD31 staining, resulting in smaller and more disconnected vascular structures. Notably, the DMSO vehicle control condition showed similar CD31 staining to the untreated control, indicating no significant impact on vascular structures. Quantitative analysis of the confocal images highlighted the differential effects of the drug treatments on vascular integrity (**Fig. 5D**). L-NNA treatment significantly increased CD31 staining intensity compared to the DMSO control, indicating enhanced vascularization. However, there is also reorganization into clustered vascular structures, as observed in the confocal images. In contrast, there is no significant difference in CD31 intensity between CA-4 treated cuboids and either control condition. The percentage of CD31 staining volume was significantly lower in CA-4 treated cuboids compared to the DMSO control, while L-NNA showed no significant difference in CD31 staining volume compared to controls. Branch length analysis revealed no difference between the two control conditions, but L-NNA treatment resulted in significantly longer total branch length, whereas CA-4 led to shorter branches. L-NNA increased the average diameter of vascular branches compared to the DMSO control, while CA-4-treated cuboids had smaller diameters, though not significantly different from the controls (**Fig. 5E**). Representative examples of 3D visualizations of CD-31 immunostained cuboids exposed to L-NNA, CA-4, and DMSO can be visualized in **Movie S**1. These quantitative measures underline the distinct vascular effects of L-NNA and CA-4, with L-NNA supporting the reorganization of the vascular structure while CA-4 impairs it.

To counteract the effects of vascular disruption, we explored the potential of nitroglycerin (NTG) to rescue and support vascular structures. NTG acts as a NO donor and is metabolized within the vascular endothelium to release NO. The process involves enzymatic activity, particularly the involvement of mitochondrial aldehyde dehydrogenase, which converts NTG into NO. The exogenous NO promotes vasodilation and oxygen delivery to tissues.^45,46^ The effect is crucial in supporting endothelial cell health and may contribute to the overall structure of the microvascular network. In a separate experiment, we assessed the effects of NTG treatment on vascular structures in cuboids. Confocal images (**Fig. 5F**) show consistent CD31 staining across all conditions, indicating stable vascular structures. Quantitative analysis (**Fig. 5G**) reveals that CD31 average intensity is significantly higher in cuboids treated with NTG at 1 µM and 50 nM under flow conditions compared to the control at 0.1 mL/hr. The 50 nM NTG no-flow condition shows a slightly higher average intensity than the control, though the difference is not significant. The CD31 staining volume percentage is also higher in cuboids treated with NTG at 1 µM and 50 nM underflow, with significant differences observed only for the 50 nM flow condition. Similarly, NTG at 50 nM without flow shows a slight, non-significant increase in volume compared to the control. For branch length, NTG treatments at 1 µM and 50 nM under flow conditions result in longer branches, with significant differences observed only for the 50 nM flow condition. The NTG 50 nM no-flow condition shows slightly longer branches than the control, but this difference is also non-significant. **Fig. 5G** further supports these findings, showing that NTG treatments at 1 µM and 50 nM under flow conditions significantly increase average branch diameter compared to the control, with no significant differences observed between the NTG 50 nM no-flow condition and the control. Representative examples of 3D visualizations of CD-31 immunostained cuboids exposed to NTG and DMSO can be visualized in **Movie S**2. Overall, these findings indicate that NTG treatment under flow conditions improves CD31 intensity, staining volume, and branch length in our microfluidically perfused cuboids. The outcome highlights the role of controlled perfusion, combined with the biomechanical effects of NTG, in maintaining and enhancing microvascular integrity in cultured tissues. The combination of biochemical treatment and mechanical flow effects demonstrates the importance of an integrated approach for optimal vascularization in tissue culture.

## 3. DISCUSSION

Our study highlights the role of fluid flow in supporting microvascular structures within cuboidal μDTs (“cuboids”) with an intact TME. Given the large variability in the biological outcome measured in this study (microvasculature architecture), our microfluidic platform for cuboids offers a convenient experimental route to obtain averages of many data points. The main limitation of our approach is that, being based on a terminal CD31 staining assay, it cannot discern between microvasculature that is *generated de novo* versus microvasculature that is *maintained*. Nevertheless, our findings demonstrate that control of the generation and/or maintenance of microvasculature within μDTs requires defined fluid flow control. These conditions are crucial as they likely influence key mechanisms such as regulating endothelial cell function, vascular stability, and efficient nutrient delivery, which are important for microvascular structure.

Our experiments with nitrogen oxide (NO) modulators indicate that flow rates between 0.05 and 0.15 mL/hr optimize the conditions that are favorable to supporting healthy microvascular environments, ^58,72^ leading to enhanced microvascular characteristics, such as increased stability and/or improved vasculature formation. This improvement is likely due to the combined effects of mechanical perfusion, NO-mediated vasodilation, and interstitial flow, which enhance endothelial cell survival and reduce apoptosis, thus contributing to the support of endothelial barrier function.^21,50,57^ Conversely, flow rates outside this optimal range, such as 0.02 mL/hr or 0.2 mL/hr, fail to support robust microvascular structures. Additionally, both orbital shaking and static conditions lacked controlled directional flow and did not achieve the same levels of observed vascular structure or molecular penetration. These findings emphasize the necessity of active perfusion and underscore the importance of maintaining appropriate flow rate thresholds for vascular support. It is also worth noting that the flow rate applied here is the bulk flow rate at the main outlet, divided among 96 traps, meaning each cuboid experiences a proportionate flow rate. Future work could explore optimizing this distribution further to enhance consistent perfusion throughout the tissue.

Our L-NNA experiments indicate that L-NNA treatment enhances CD31 intensity and branch length, suggesting increased endothelial cell density and vascular network complexity. We also observed a significantly higher branch average diameter of vasculature under L-NNA treatment. The increase in branch diameter may be a compensatory response to maintain blood flow in the face of increased vascular resistance. However, the reorganization of vascular structures into clustered formations implies that NO inhibition disrupts normal vascular morphology, potentially a compensatory mechanism for reduced NO availability, leading to a more compact and possibly less functional vascular network. Thus, using L-NNA to inhibit NOS and reduce NO production provides valuable insights into the complex role of NO in vascular biology. While it enhances certain aspects of vascular structure, it also disrupts normal vascular morphology, highlighting the balance NO plays in structuring a functional and adaptive vascular network.

CA-4 treatment resulted in reduced CD31 staining intensity within the cuboids, indicating endothelial cell loss and vascular network disruption. The microvascular structures within the cuboids fragmented, a direct consequence of microtubule destabilization, which led to endothelial barrier breakdown and vessel collapse. The vascular shutdown causes tumor hypoxia and nutrient deprivation, ultimately leading to tumor cell death and necrosis.^44^ These results highlight CA-4’s therapeutic potential in targeting tumor vasculature by destabilizing microtubules and disrupting the microvascular network.

Our results indicate that NTG’s effects are concentration-dependent.^54^ At lower concentrations, NTG appears to support vascular structures even without flow, likely due to a sufficiently basal level of NO, promoting endothelial cell survival and preventing vascular collapse. However, the effects are less pronounced at higher concentrations, suggesting an optimal concentration range for NTG in therapeutic applications. The combination of biochemical stimuli from NTG and mechanical stimuli from controlled fluid flow synergistically influence vascular characteristics. NTG provides a consistent NO supply, promoting vasodilation and endothelial cell function, while controlled fluid flow mimics physiological shear stress, potentially activating eNOS and endogenous NO production.^58^ Together, these stimuli contribute to a more robust and stable vascular network, with significant implications for tissue engineering and therapeutic applications.^59^

Despite the promising results, our study has several limitations. While the microfluidic device supports some vascularization and facilitates controlled perfusion, it still falls short of fully replicating the complexity of the *in vivo* TME and its highly dynamic and interacting metabolic and immunological states. Structurally, the device does not effectively interface with the biological vasculature, particularly in forming anastomoses, which are essential for establishing direct blood flow connections between the engineered microvasculature and the host tissue’s vascular network. Incorporating endothelial cells outside the cuboid to establish connections with the vasculature within the cuboid could improve the model by creating a more realistic and functional vascular network.^33,34,60–64^ Potential variability in cuboid size and shape may introduce inconsistencies in perfusion and drug delivery despite our normalization efforts. Moreover, while we thoroughly investigated the effects of different drugs, the long-term viability and stability of vascular structures under prolonged treatment conditions remain to be fully established, requiring further studies to evaluate chronic exposure effects and potential sustained therapeutic benefits. Addressing these limitations in future research is crucial for advancing the utility and accuracy of *in vitro* tumor models.

Additionally, future research should focus on refining flow conditions and exploring the long-term effects of various flow rates and drug treatments on tumor vasculature. Establishing long-term flow (beyond 72 hrs) is crucial, and systematic studies are needed to determine the optimal duration and intensity of perfusion required to sustain microvascular structures over extended periods, including continuous versus intermittent flow, varying shear stress levels, and different perfusion patterns. Our results indicate that while current flow conditions support microvascular structures more effectively than static conditions at 72 hrs, there is a notable decline in vascular structures compared to the 24-hr mark. This finding suggests that additional or adjusted dynamic flow (*e.g.*, peristaltic) regimens may be necessary to sustain the perfusion of microvasculature over extended periods. Furthermore, while our study primarily focuses on the vascular aspect of the TME, further research is needed to explore how vascular structures interact with other cellular elements within the tumor, such as immune cells and fibroblasts.^65^ Future studies should investigate the effects of these complex multicellular components that could provide more in-depth insights into TME dynamics and therapeutic responses, potentially leading to more effective immunotherapies and stromal-targeted treatments. While our current study offers valuable insights into controlled perfusion for supporting microvascular structures within tumor cuboids, future research should aim to expand the capabilities of *in vitro* tumor models, ultimately improving cancer research and treatment outcomes.

## 4. METHODS AND MATERIALS

### 4.1. Reagents and Materials

All reagents were obtained from Thermo Fisher Scientific (MA, USA) unless otherwise specified. Dextran Texas Red (70,000 MW, Lysine Fixable) was procured from Invitrogen (Waltham, MA). 5-(N-2,3-Dihydroxypropylacetimido)-2,4,6-triiodo-N, N’-bis-(2,3dihydroxypropyl) isophthalamide (Histodez) was purchased from MP Biomedical (Santa Ana, CA). N-Methylacetamide (99.0%) and Alpha-Thioglycerol (98.0%) were obtained from TCI America (Portland, OR). Normal Mouse Serum was purchased from Life Technologies Corp (Carlsbad, CA). Alexa Flour 488 anti-mouse CD31 Antibody was purchased from BioLegend (San Diego, CA). Deionized water (DI) was purified using Milli-Q (Millipore, Bedford, MA). Drugs were diluted from DMSO stocks (10–200 mM), purchased from MedChem Express, Nω-Nitro-L-arginine (L-NNA, N5751), Combretastatin A4 (CA-4, C7744), Trinitroglycerin solution, 1% (w/w) in propylene glycol (T-021) were purchased from Millipore Sigma (Burlington, MA). PMMA was purchased from Astra Products, Inc. (NY, USA). Detailed storage conditions, handling precautions, and reagent preparation procedures are outlined in the subsequent sections.

### 4.2. Design and Fabrication of a Multi-layer Microfluidic Device

We utilized AutoCAD 2021 (Autodesk, CA, USA) to design the layers of the device. Our microfluidic platform features three operational layers: a sealing layer, a fluidic layer, and a loading frame. We micromachined PMMA using a CO_2_ laser system, adjusting laser powers and speeds for optimal fabrication of each layer. The CO_2_ laser system (VLS3.60, Scottsdale, USA) we operated has a wavelength of 10.6 μm, and a maximum power of 30 W. We imported all 2D files from AutoCAD into the Universal Control Panel (UCP) software of the VLS3.60 to proceed with the manufacturing. We performed a raster cut on 0.8 mm thick PMMA for the microchannel layer, shaping channels ranging from 400 to 500 µm in width and 250 to 350 µm in depth. We designed cylindrical cuboid traps with an 800 µm depth and 600 µm diameter to accommodate a single cuboid each. We precisely cut the sealing layer from a 0.3 mm thick PMMA. To serve as a reservoir for the liquid suspension of cuboids, we fabricated a loading frame with a thicker PMMA of 6.4 mm. We manually applied 3M™ High-Strength Acrylic Adhesive 300LSE (MN, USA) to line the loading frame before cutting. We cut through-holes into the loading frame’s perimeter and inserted silicone tubes (1/16“ ID, 1/38” OD; Cole Parmer) as microfluidic inlets. Small reservoirs created by raster cuts positioned near the inlets facilitated the accumulation of cyanoacrylate glue, ensuring a durable tube attachment when assembling the device.

Following laser ablation of PMMA, which generates polymer debris and reflow, we implemented an established cleaning and assembly protocol.^32^ Specifically, we rinsed the laser-cut components with DI water. This step was followed by a 30-second sonication in an isopropanol (IPA) bath, effectively removing any remaining debris. We exposed the channel networks to chloroform vapor to improve optical clarity and reduce surface roughness.^66^ We conducted this in a glass container (264 mm (L) × 213 mm (W) × 165 (T) mm) with 50 mL of chloroform at the base. By placing the components on steel standoffs 6 mm high, we maintained a 3 mm gap above the chloroform. Both the channel network and sealing layers underwent a 5-minute chloroform vapor exposure.

Immediately following the chloroform vapor treatment, we began the bonding process using a combination of thermal and solvent techniques. Exposure to chloroform vapor renders the PMMA surface slightly sticky due to the reflow of the polymer.^66^ The PMMA surfaces softened from the solvent and achieved a molecular bond as they came into contact with the remaining vapor dispersed from their juncture. We first pressed the layers together by hand to establish an initial bond. To secure a consistent bond, we positioned the pair between two PDMS slabs, approximately 3 mm-thick and matching the channel layer in size. We then processed the assembly in a heat press (Model 3912, Carver, IN, USA) at 200 psi and 60 °C for 4 minutes. To assemble the loading frame before cuboids loading, we peeled off the 3M300LSE liner, manually pressed the components together, and then applied the heat press again for five minutes at 200 psi to guarantee a leak-free bond.

### 4.3. Tumor Generation for Mouse Model

Mice were managed following institutional guidelines and under protocols endorsed by the Animal Care and Use Committee at the Fred Hutchinson Cancer Research Center (Seattle, USA). Mouse syngeneic Py8119 breast cancer cells suspended at 1–2 million/150 μL of 33% (v/v) Matrigel (Thermo Fisher Scientific) were orthotopically injected into 6-8-week-old female C57BL mice (Jackson Laboratories). The mice were euthanized before the tumor volume exceeded 2 cm^3^ (within 2–4 weeks).

### 4.4. Cuboid Preparation

Following an established protocol, we generated the cuboidal-shaped microdissected tissues using a previous protocol.^32^ In brief, we first dissected and discarded the necrotic regions. We then embedded tissue punches (600 μm diameter) in 2% low-melting agarose. Using a 5100 mz vibratome (Lafayette, Instrument), we sliced the tissue into 400 μm-thick slices in an ice-cold HBSS solution (1X) and transferred the slices into ice-cold DMEM-F12 with 15 mM HEPES (Invitrogen). We cut the slices into cuboids using a standard Mcllwain tissue chopper (Ted Pella, Inc., USA) by making two orthogonal sets of cuts and rotating the sample holder between sets. Next, we separated the cuboids by gently pipetting them up and down in DMEM-F12-HEPES and filtered them through 750 μm and 300 μm mesh filters (Pluriselect, USA). We performed three washes using DPBS and concluded with a final rinse in DMEM-F12 containing HEPES. The prepared cuboids were subsequently maintained on ice until they were ready for loading.

### 4.5. Device Preparation before Cuboid Loading

To prepare for the loading of live tumor tissues, we treated each device with oxygen plasma for 5 min at approximately 950 mTorr and 60 W using a Diener RF plasma oven, which increased the hydrophilicity of the PMMA surfaces. Then, to ready the device for its application, we manually filled it with 100% ethanol using 60 mL syringe (BD Bioscience, San Jose, CA) to displace any air within the trapping regions. After we ensured the complete removal of bubbles, we flushed the microchannels first with sterile DI water and then with sterile PBS. We further sterilized the devices by cycling them with 70% ethanol, sterile DI water, and sterile PBS twice before loading and culturing cuboids.

### 4.6. Cuboids Loading Process

We began by suspending the cuboids in a 20% w/w PEG (8,000 M.W.) in PBS solution within a Corning Falcon round-bottom polystyrene tube. We filled the open-top area of the device with the same PEG solution and used a 60 mL syringe on a syringe pump (Fusion 200, Chemyx Inc., Stafford, TX) to pull the solution into the microchannels. Dispensing multiple cuboids into the device’s open top with a 3 mL transfer pipette, we observed that most cuboids floated above the traps, with a minority settling inside. We then initiated hydrodynamic suction by setting the syringe pump to a flow rate of 200 mL/hr, ensuring efficient trapping of the cuboids without rapid depletion of the PEG solution. Keeping the PEG/PBS solution cold, we continuously refill the open-top area with fresh PEG solution, amounting to approximately 50 mL. The cuboids needed to be close to the trap (within approximately 400 μm) for the suction to capture them effectively. We used tweezers to carefully position the cuboids near the activated traps, ensuring precise loading of one cuboid per trap. By employing this technique, we successfully loaded cuboids into the quadrant network activated by suction until all traps were filled. Once we had filled the network with cuboids, we aspirated any excess PEG and replaced it with Dulbecco’s Phosphate-Buffered Saline (DPBS), then aspirated again, finally introducing a culture medium. The medium contained Dulbecco’s Modified Eagle Medium: Nutrient Mixture F12 (DMEM-F12, Thermo Fisher), 10% heat-inactivated fetal bovine serum (FBS, VWR), and 1% penicillin/streptomycin (P/S, Invitrogen).

### 4.7. Cuboid Culture in Device with Perfusion

After loading the cuboids into the devices and introducing the culture medium, we placed the devices in a cell culture incubator for controlled perfusion. We connected syringe pumps to the devices’ outlet tubing in withdraw mode, setting flow rates ranging from 0.02 mL/hr to 0.2 mL/hr using 30 mL syringes. Alongside these, we maintained one set of control devices with orbital shaking but no-flow, and another set was completely static without shaking. We collected and assessed the devices under these different conditions at 2 hrs, 24 hrs, and 72 hrs intervals. For the experiments spanning 24 hrs and 72 hrs, we refilled the culture medium every 12 hrs to keep the solution level above the traps, thereby preserving tissue viability and ensuring a steady supply of nutrients.

### 4.8. Dextran Perfusion Assessment

To assess the effectiveness of our perfusion system, we introduced dextran Texas Red 70 kDa into the culture medium to a final concentration of 10 mg/mL. We replaced the standard culture medium with this dextran-containing medium and cultured the devices in the incubator under the previously established conditions, maintaining perfusion and control setups. To monitor the progress of the perfusion, we conducted the dextran perfusion assessment over two periods, 2 hrs and 24 hrs. For the 24-hr duration, we refilled the dextran solution at the 12-hr mark to ensure a constant supply throughout the experiment.

### 4.9. Cuboid Culture with Perfusion and Drug Administration

Following the previously outlined protocol, we cultured cuboids in our microfluidic devices, administering pharmaceutical agents at a steady flow rate of 0.1 mL/hr. For one set of experiments, we administered L-Nω-Nitro arginine methyl ester (L-NNA) at 10 μM and Combretastatin A-4 (CA-4) at 10 μM, both diluted in the culture medium containing a 0.1% v/v DMSO stock solution. We benchmarked these treatments against control groups: one received only a 0.1% v/v DMSO vehicle at the same flow rate, and another was subject to the same flow rate but without DMSO, each over 24 hrs. Separately, we exposed devices to nitroglycerin (NTG) at concentrations of 50 nM and 1 µM,^54^ which did not require dilution in DMSO, maintaining the 0.1 mL/hr flow rate. A no-flow culture device also received 50 nM nitroglycerin treatment. The control for this setup included a flow without drug presence for 24 hrs. We replenished the culture medium in each device halfway through at 12 hrs. Following the 24-hr incubation, we retrieved the devices for subsequent analysis. All pharmaceutical agents were stored at -20 °C upon receipt.

### 4.10. Tissue Preparation and Immunostaining

We conducted whole-mount immunostaining within the microfluidic devices using an adapted protocol from Li et al. (2019).^67^ Initially, we fixed the samples in 4% (v/v) paraformaldehyde (PFA) in PBS, allowing them to remain overnight at 4 °C. Subsequently, the fixed tissues were washed extensively with PBS for 30 to 60 min, repeated with two additional washes. For blocking and permeabilization, we incubated them at 37 °C for 8 to 24 hrs on a shaker. We used a freshly prepared conventional blocking buffer composed of 1× PBS with 0.3% Triton X-100, 1% BSA, and 1% normal mouse serum, which was passed through a 0.45-μm filter to minimize bacterial contamination. For staining, we incubated the tissues in a blocking buffer containing Alexa 488 anti-mouse CD31 (1:250, BioLegend) for 3 days in a shaking incubator set at 37 °C and 150-220 rpm. We ensured the microfluidic devices were sealed with Parafilm to prevent evaporation and protected from light to prevent photo-degradation. After the staining incubation, we washed the tissues with a washing buffer of 1x PBS solution supplemented with 0.3% Triton X-100 and 0.5% 1-thioglycerol at 37 °C for 8-14 hrs, careful not to extend washing and risk destabilizing the staining or losing the signal. We then refreshed the tissues with a new washing buffer. The devices can sit at room temperature for 1-4 days, refreshing the buffer every 12-24 hrs before clearing.

### 4.11. Preparing Clearing Solution and Tissue Clarification

To prepare the Ce3D clearing solution, we first incubated a bottle of 100% N-methyl acetamide at 37 °C for over an hour to liquefy it since it is solid at room temperature. To facilitate its transfer, we pre-warmed a 25-mL pipette before drawing 20 mL of the N-methylacetamide to avoid adherence to the pipette walls. We transferred the solution to a 50-mL conical tube to prepare a 40% N-methyl acetamide stock solution by adding 30 mL of room-temperature PBS and mixing thoroughly. For the clearing mixture, 16 g of Histodenz was layered between 8 mL and 3 mL of the 40% N-dimethyl acetamide solution in a 50-mL conical tube and 20 µL of Triton X-100. The Histodenz dissolves more efficiently when encapsulated within the N-methyl acetamide layers. After sealing the tube with Parafilm, we placed it in a 37 °C shaking incubator at 150–225 rpm until the contents fully dissolved. To the final Ce3D clearing solution, we added 100 µL of 1-thioglycerol to prevent discoloration. Before adding the clearing solution to the device, we blotted the samples using Kimwipe to remove excess washing buffer. We wrapped the device in aluminum foil to protect it from light. Finally, the device was placed on a shaker at room temperature for 24 hrs to complete the tissue clarification.

### 4.12. Fluorescence Imaging Microscopy

We employed a Leica TCS SP8 confocal microscope (Leica Microsystems, Germany) for high-resolution imaging of cleared tissues. The instrument was set to a scan speed of 600 Hz, and image resolution was formatted to 1024 x 1024 pixels. We employed a 10x air objective lens and adjusted the pinhole to 1 Airy Unit (approximately 38.4 µm), optimizing the spatial resolution. We acquired a z-series stack for each cuboid to enable three-dimensional reconstruction with a z-step interval of 2 µm. Two optically pumped semiconductor lasers (OPSLs) provided illumination at wavelengths of 488 nm and 552 nm, respectively. The 488 nm laser was used to excite Alexa 488 conjugated to CD31, while the 552 nm laser specifically targeted dextran Texas Red. Hybrid detectors (HyD) were utilized for detection due to their enhanced sensitivity and broader dynamic range, ensuring clear and distinct visualization of each fluorophore.

### 4.13. Image Processing and 3D Reconstruction

We processed the volumetric images of cuboids using Imaris software (9.9.0 Bitplane, UK) to perform 3D rendering and visualization. We utilized the Surface tool within Imaris for vascular reconstructions. We enhanced precision in delineation by creating CD31 surfaces with Labkit (Image J, NIH), incorporating machine learning for advanced segmentation.^68^ We trained a classifier to identify vascular features from a representative stack of cuboid images across all experimental conditions for each time point of 2 hrs, 24 hrs, and 72 hrs. We then consistently applied this classifier to the entire set of images within each experimental time point. To improve the clarity and definition of the CD31 channel, we applied a gaussian filter and background subtraction before creating the surfaces. Additionally, we utilized CD31-labeled surfaces as masks to generate detailed filament structures, enhancing our quantitative analysis of the microvasculature network. The branch length was measured using the Imaris filament tracing function, which calculates the total length of branches extending from the vasculature’s main body within the cuboids. Similarly, the branch average diameter was calculated by averaging the local diameters measured at specific points along each branch, also using the Imaris filament tracing function. For cuboids undergoing dextran perfusion, we reconstructed their outer surfaces to evaluate the dextran’s penetration depth and intensity distribution, facilitating comparative infiltration analysis across various experimental conditions.

### 4.14. Statistical Methodology

We utilized GraphPad Prism 10 (Boston, MA, USA) to conduct statistical analyses and generate data visualizations. Additionally, we employed MATLAB (R2021a, USA) to further analyze and export experimental data from Imaris. The selection of statistical tests was based on preliminary assessments of normality, which showed whether the data conformed to a normal distribution. If the data passed the normality test, we used one-way ANOVA with Tukey’s post-test. If the data did not pass the normality test, we applied the non-parametric one-way ANOVA, Kruskal-Wallis test, followed by Dunn’s multiple comparison test to compare multiple experimental groups against the control group. We denoted significance levels as *** for p<0.001, ** for p<0.01, * for p<0.05, and ns (not significant) for p>0.05. Non-parametric tests are advantageous in cases where the data do not conform to a normal distribution as they do not assume normal distribution, thereby providing a suitable alternative for our dataset. Their application ensured the robustness of our statistical analysis against the non-normality of the data.

## Supporting Information

Supporting Information is available from the Wiley Online Library or from the author.

## Acknowledgements

This work was supported by the National Cancer Institute of the National Institutes of Health (R01CA181445-A.F, R01CA272677-A.F), and Brotman Baty Institute, 2021 Catalytic Collaborations Trainee Award (T.N.H.N).

## Supplementary Materials

**Fig. S1.** Correlation of cuboid volume with normalized dextran intensity under different flow rates and no-flow control.

**Fig. S2.** Analyze cuboid volume cut-off on dextran average intensity under different flow rates and no-flow control.

**Fig. S3** Correlation of cuboid outer volume with the percentage of CD31 staining volume under different flow rates and no-flow control for 2 hrs, 24 hrs, and 72 hrs.

**Movie S1.** Visualization of cuboids corresponding to Fig. 5D-E.

**Movie S2:** Visualization of cuboids corresponding to Fig. 5F-G

