## Supplementary Materials for "Microfluidic Modulation of Microvasculature in Microdissected Tumors"


**This PDF file includes:**

Figs. S1 to S3

**Other Supplementary Materials for this manuscript include the following:**

Movies S1 to S2

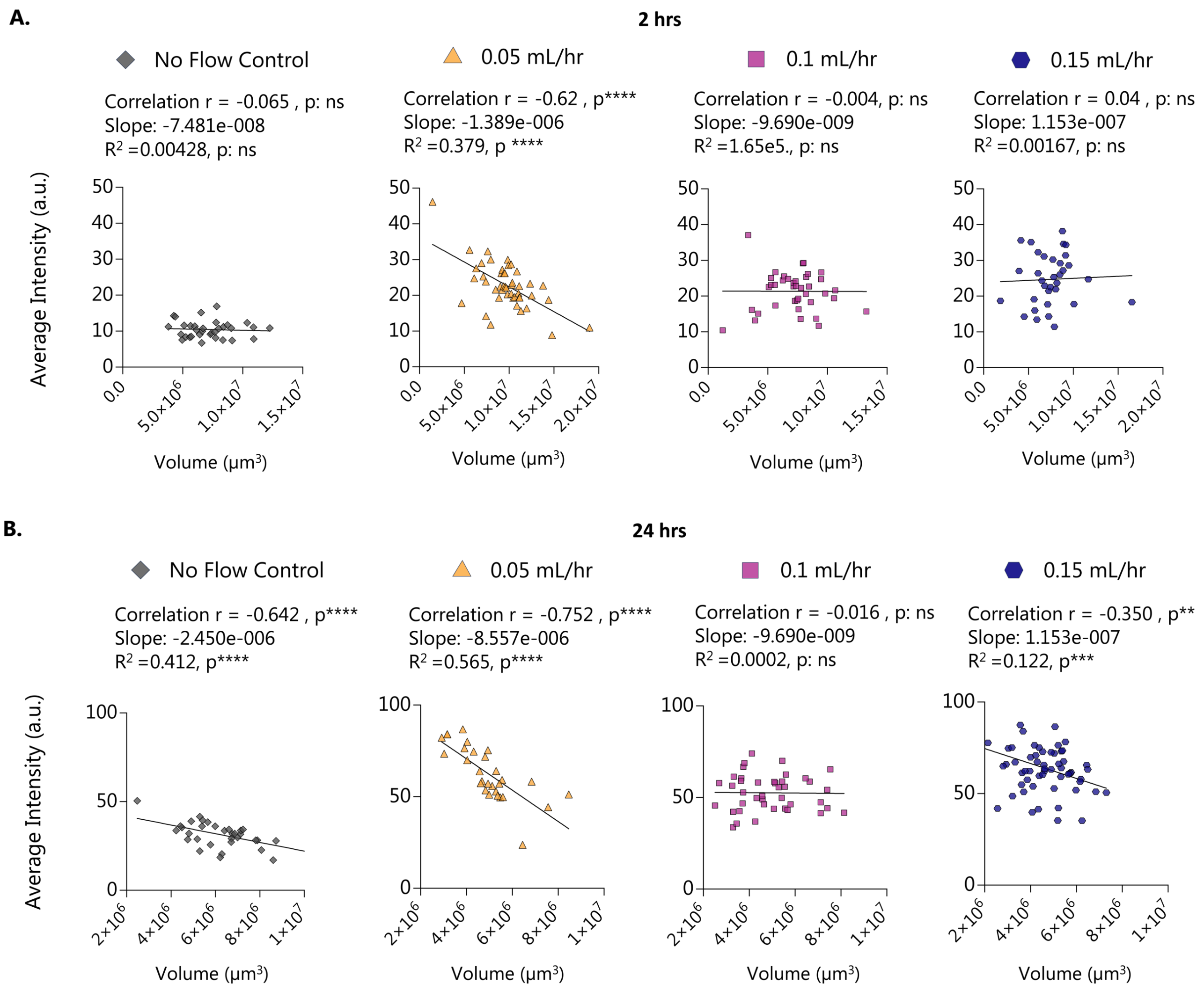


**Fig. S1. Correlation of cuboid volume with average dextran intensity under different flow rates and no flow control. A.** The relationship between cuboid outer volume and average dextran intensity at 2 hrs for various flow conditions. Each graph individually shows data for no flow conditions, 0.05 mL/hr, 0.1 mL/hr, and 0.15 mL/hr flow rates, respectively. Each graph includes Pearson correlation analysis results (r and p-value) and a simple linear regression line with corresponding slope, $R^2$ and p-values. **B.** The relationship between cuboid volume and normalized dextran intensity at 24 hrs under the same conditions. Each graph individually shows data for no flow conditions, 0.05 mL/hr, 0.1 mL/hr, and 0.15 mL/hr flow rates, respectively. Each graph includes Pearson correlation analysis results (r and p-value) and a simple linear regression line with corresponding slope, $R^2$ and p-values. *$p<0.05$, ***$p<0.001$, and ****$p<0.0001$; and 'ns' for $p>0.05$.

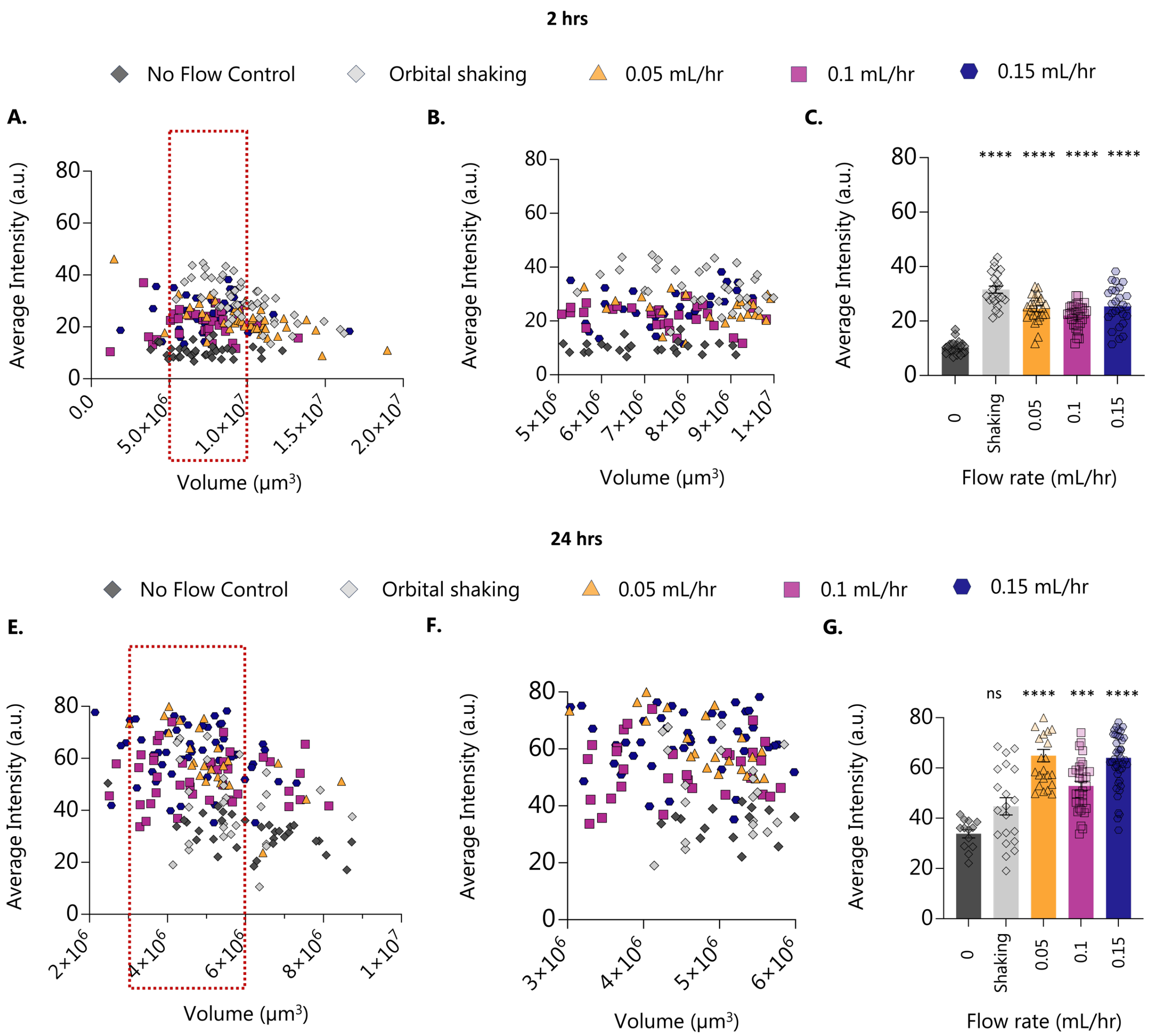


**Fig S2. Analyze cuboid volume cut-off on average dextran intensity under different flow rates and no-flow control. A.** For the 2 hrs condition, the graph shows the relationship between cuboid volume and average dextran intensity for all conditions, with a dashed box indicating the cut-off volume range, where no significant correlation is observed between cuboid volume and dextran intensity. **B.** The graph illustrates the relationship between cuboid volume and average dextran intensity, focusing on the cut-off volume. **C.** The bar graph presents the mean dextran intensity for flow rates of 0.05, 0.1, and 0,15 mL/hr compared to no-flow conditions, corresponding to the cuboid with the volume cut-off shown in B. **D.** For the 24 hrs condition, the graph illustrates the relationship between cuboid volume and average dextran intensity, with a dashed box indicating the cut-off volume range, where no significant correlation is observed between cuboid volume and dextran intensity. **E.** The graph shows the relationship between cuboid volume and average dextran intensity, focusing on the cut-off volume for all conditions where no correlation was observed between cuboid volume and dextran intensity. **F.** The bar graph presents the average dextran intensity for flow rates 0.05, 0.1, and 0.15 mL/hr compared to the no flow condition, corresponding to the cuboid with the volume cut-off shown in F. Statistical significance was determined using nonparametric Kruskal-Wallis one-way ANOVA with Dunn's post-test, comparing all conditions to no flow control with significance denoted as *$p<0.05$, ***$p<0.001$, and ***$p<0.0001$; non-significant findings are indicated by 'ns' for $p>0.05$.

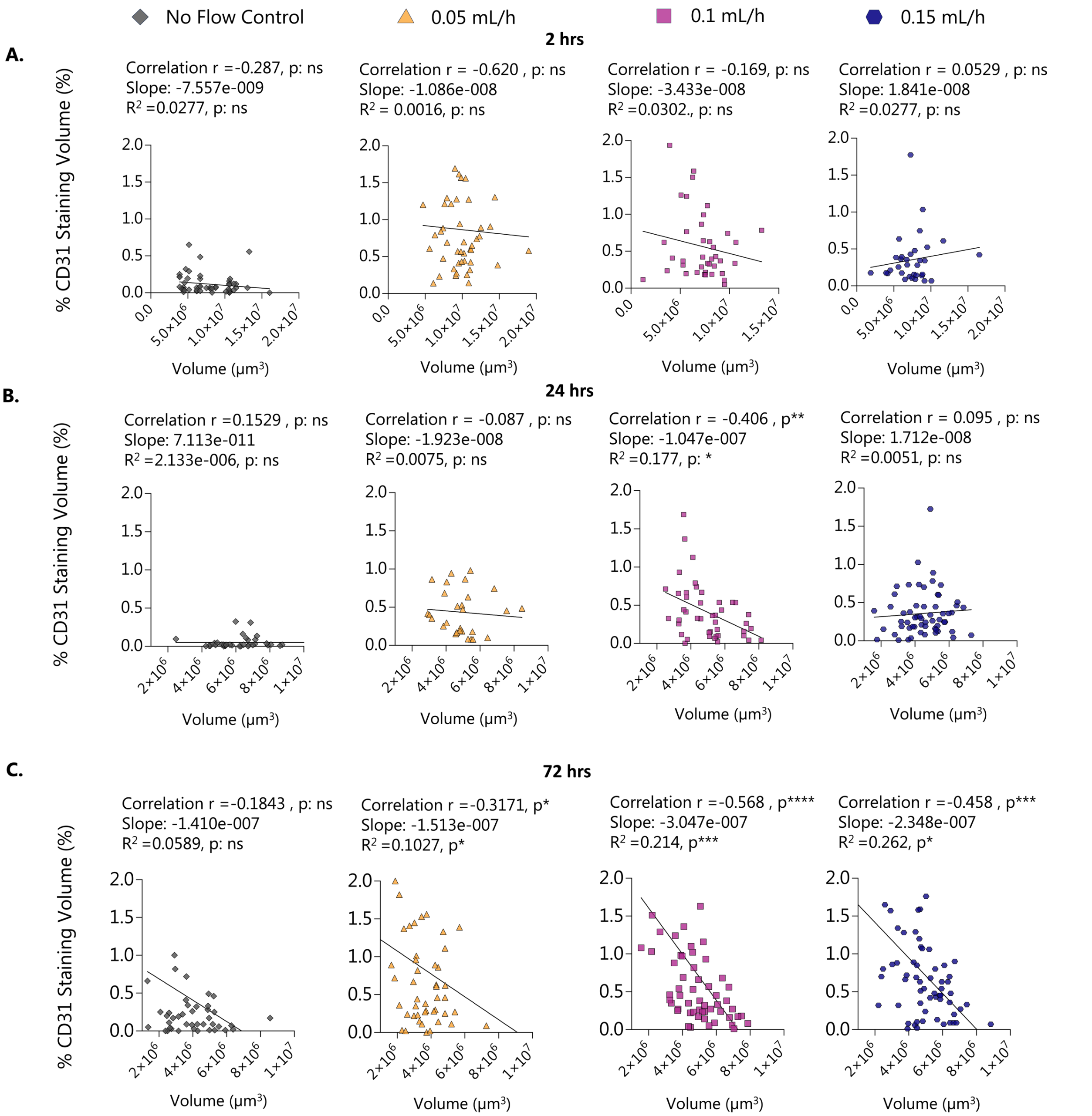


**Fig. S3. Correlation of cuboid outer volume with the percentage of CD31 staining volume under different flow rates and no flow control for 2 hrs, 24 hrs, and 72 hrs. A.** The relationship between cuboid volume and the percentage of CD31 staining surface at 2 hrs for various flow conditions. Each graph individually shows data for no flow condition, 0.05 mL/hr, 0.1 mL/hr, and 0.15 mL/hr flow rates, respectively. Each graph includes Spearman correlation analysis results (r and p-value) and a simple linear regression line with corresponding slope, $R^2$ and p-values. **B.** The relationship between cuboid volume and the percentage of CD31 staining surface at 24 hrs for various flow conditions. Each graph individually shows data for no flow conditions, 0.05 mL/hr, 0.1 mL/hr, and 0.15 mL/hr flow rates, respectively. Each graph includes Spearman correlation analysis results (r and p-value) and a simple linear regression line with corresponding slope, $R^2$ and p-values. **C.** The relationship between cuboid volume and the percentage of CD31 staining surface at 72 hrs for various flow conditions. Each graph individually shows data for no flow conditions, 0.05 mL/hr, 0.1 mL/hr, and 0.15 mL/hr flow rates, respectively. Each graph includes Spearman correlation analysis results (r and p-value) and a simple linear regression line with corresponding slope, $R^2$ and p-values. Significance denoted as *p<0.05, ***p<0.001, and ***p<0.0001; non-significant findings are indicated by 'ns' for p>0.05.

**Movie S1. Visualization of cuboids corresponding to Fig. 5D-E.** The movie shows cuboids under treatment with L-NNA, CA-4, and DMSO control vehicle. For each treatment condition, three cuboids are presented, corresponding to the high, average, and low percentages of CD31 staining volumes. This movie illustrates the differential effects of L-NNA and CA-4 treatments on microvascular structure preservation within the cuboids compared to the DMSO control.

**Movie S2: Visualization of cuboids corresponding to Fig. 5F-G.** The movie shows cuboids under different treatment conditions: 0.1 mL/hr flow with NTG 1 µM, 0.1 mL/hr flow with NTG 50 nM, and no flow with NTG 50 nM, compared to the control (0.1 mL/hr flow without any drug). Three cuboids are presented for each condition, corresponding to the high, average, and low percentages of CD31 staining volumes. This movie illustrates the effects of NTG treatment under various flow conditions on microvascular structure preservation within the cuboids.
